# Enhancing GABAergic Tone in the Rostral Nucleus of the Solitary Tract Reconfigures Sensorimotor Neural Activity

**DOI:** 10.1101/2020.02.20.958199

**Authors:** Joshua D. Sammons, Caroline E. Bass, Jonathan D. Victor, Patricia M. Di Lorenzo

**Author notes:** To whom correspondence should be addressed Dept. of Psychology, Box 6000, Binghamton University, Binghamton. NY 13902-6000. J.D. Sammons’ current affiliation is Department of Biochemistry and Molecular Genetics, University of Alabama at Birmingham, Birmingham, AL 35294-2170.

## Abstract

Recent work has shown that most cells in the rostral, gustatory portion of the nucleus tractus solitarius (rNTS) in awake, freely licking rats show lick-related firing. However, the relationship between taste-related and lick-related activity in rNTS remains unclear. Here, we tested if GABA-derived inhibitory activity regulates the balance of lick- and taste-driven neuronal activity. Combinatorial viral tools were used to restrict expression of ChR2-EYFP to GAD1+ GABAergic neurons. Viral infusions were bilateral in rNTS. 2-4wks later, an optical fiber attached to 8-16 drivable microwires was implanted into the rNTS. After recovery, water-deprived rats were presented with taste stimuli in an experimental chamber. Trials were 5 consecutive taste licks [NaCl, KCl, NH_4_Cl, sucrose, MSG/IMP, citric acid, quinine, or artificial saliva (AS)] separated by 5 AS licks on a VR5 schedule. Each taste lick triggered a 1s train of laser light (25Hz; 473nm; 8-10mW) in a random half of the trials. In all, 113 cells were recorded in the rNTS, 50 responded to one or more taste stimuli without GABA enhancement. Selective changes in response magnitude (spike count) within cells shifted across unit patterns but preserved inter-stimulus relationships. Cells where enhanced GABAergic tone increased lick coherence conveyed more information distinguishing basic taste qualities and different salts than other cells. In addition, GABA activation significantly amplified the amount of information that discriminated palatable vs. unpalatable tastants. By dynamically regulating lick coherence and remodeling the across-unit response patterns to taste, enhancing GABAergic tone in rNTS reconfigures the neural activity reflecting sensation and movement.

**Significance Statement:** The rostral nucleus tractus solitarius (rNTS) is the first structure in the central gustatory pathway. Electrophysiological recordings from the rNTS in awake, freely-licking animals show that cells in this area have lick- as well as taste-related activity, but the relationship between these characteristics is not well understood. Here, we showed evidence that GABA activation can dynamically regulate both of these two properties in rNTS cells to enhance the information conveyed, especially about palatable vs. unpalatable tastants. These data provide insights into the role of inhibitory activity in the rNTS.

## INTRODUCTION

In mammals, information about gustatory stimulation is conveyed directly to the rostral nucleus tractus solitarius (rNTS). This structure directs taste information to higher order structures, integrates information from centrifugal sources and, ultimately, influences movements aimed at ingestion. In rNTS of alert rats, only a minority of cells are taste-responsive; most cells, including taste-responsive cells, track behavior (Denman et al., 2019). That is, when rats freely lick tastants of various qualities, coherence of firing patterns with the lick cycle is very common (Denman et al., 2019). Moreover, lick-related cells also contribute information about taste quality along with canonically taste-responsive cells, albeit at a lower level. Cells that show global shifts in firing rates during licking have also been described (Denman et al., 2019; Roussin et al., 2012; Weiss et al., 2014), underscoring the intimate relationship of rNTS activity with behavior. Thus, an essential feature of the rNTS in awake animals is that activity reflects both taste sensation and the movements associated with ingestion.

In addition to the sensorimotor aspects of the rNTS, the extent to which taste responses in rNTS cells can be altered by a variety of physiological and experimental conditions reveals a surprising amount of plasticity. For example, taste responsivity within a cell can be altered by taste adaptation (Di Lorenzo and Lemon, 2000), differences in taste context (Di Lorenzo et al., 2003) or the simple passage of time (Sammons et al., 2016), even to the point where taste responses that were not previously evident were uncovered. Further, suppression (Monroe and Di Lorenzo, 1995) or stimulation (Smith and Li, 2000) of the gustatory cortex, lateral hypothalamus (Cho et al., 2002, 2003; Matsuo et al., 1984; Murzi et al., 1986) and amygdala (Cho et al., 2003; Li et al., 2002), all of which provide centrifugal input to rNTS, can selectively alter responses to individual tastants in rNTS cells.

One potential mechanism that may underlie or contribute to these changes is the action of GABA in the NTS, since several structures that send descending input to the rNTS either synapse on GABAergic interneurons (Smith and Li, 2000) or provide GABAergic input directly to rNTS neurons (Saha et al., 2002).

The presence of GABA in the rNTS has been well documented (Boxwell et al., 2013; Davis, 1993; Lasiter and Kachele, 1988), but the functional consequences for the neural representation of taste are still not fully understood. Leonard et al. (1999) argued that the anatomical localization of GABAergic terminals on dendrites in rNTS facilitated GABAergic modulation of incoming gustatory signals. In physiological studies, Grabauskas and Bradley (1998; 1999) showed that tetanic stimulation of the solitary tract induces both short- and long-term GABA-mediated potentiation of inhibitory synaptic activity, suggesting that this type of presynaptic plasticity may aid in stabilizing the response to afferent input (Grabauskas and Bradley, 1999). In addition to inhibition produced by afferent signals, taste-responsive cells in the rNTS are under tonic inhibitory influence (Grabauskas and Bradley, 2003; Smith and Li, 1993) presumably derived from GABAergic interneurons. Moreover, application of the GABA antagonist bicuculline can broaden the breadth of tuning of taste-responsive rNTS cells. Moreover, inhibitory interactions in rNTS may enhance and stabilize the temporal structure of taste-evoked spike trains (Rosen and Di Lorenzo, 2009). The caveat to what is known about the functionality of GABA-driven inhibition in neural coding of taste is that it is all derived from studies in anesthetized subjects; the function of inhibition in taste coding in awake subjects may be different.

Here, we used optogenetic tools to selectively enhance GABAergic activity in the rNTS while rats were freely licking taste stimuli. Results showed that GABA activation can modify taste responses in a stimulus-selective manner in a subset of cells. In some cases, GABA enhancement produced taste responses in cells that were unresponsive otherwise; in other cases, taste-responsive cells became completely unresponsive. Finally, GABA activation could modulate lick coherence and that predicted changes in taste profiles.

## MATERIALS AND METHODS

### Subjects

Nine male (250-450 g) and two female (200-350 g) Sprague-Dawley rats obtained from Taconic Laboratories (Germantown, New York) served as subjects. Food and water were provided *ad libitum* except during behavioral studies where rats were water deprived for 22-23hrs per day. Rats were pair housed and maintained on a 12 h light-dark cycle with lights on at 2100 hours. All procedures were approved by the Institutional Animal Care and Use Committee of Binghamton University and conducted in accordance with the National Institutes of Health Animal Welfare Guide.

### Viral constructs and infusion

Rats were anesthetized with a ketamine:xylazine mixture (100 mg/kg:14 mg/kg, i.p.). Buprenorphine-HCl (0.05 mg, s.c.) was administered to enhance the effects of the anesthetic and atropine sulfate (0.054 mg/kg, s.c.) to prevent excessive secretions. The rat’s scalp was shaved and its head was secured in a stereotaxic instrument (David Kopf Instruments, Tujunga, CA). The head was leveled with bregma and lambda in the same dorsal-ventral plane. The rat’s eyes were lubricated and core temperature maintained at 37 °C with a heating pad attached to an anal thermistor probe. The scalp was then swabbed three times with Betadine alternated with 70% ethanol. An incision was made along the midline from bregma to the occipital ridge and the skin and fascia were retracted with blunt dissection. A hole was drilled at 12 mm posterior and ±1.75 mm lateral to bregma. A combination of viruses was infused (0.5 µL total; 0.5 µL/min) bilaterally 6 mm below the surface of the brain. The combination consisted of 166 nL of GAD1-Cre-AAV 2/10 + 333 nL of Ef1α-DIO-ChR2-EYFP-AAV 2/10, which we have previously shown to restrict expression to GAD1+ neurons (Xiao et al., 1998). All viruses were packaged using the triple transfection method to generate pseudotyped virus as detailed elsewhere (Gompf et al. 2015). After each infusion, the needle was held in place for an additional 5 minutes to ensure complete expulsion of the virus. After retraction of the needle, the scalp was sutured and the rat allowed to regain consciousness. The animal was given a post-operative injection of buprenorphine-HCl (0.05 mg; s.c.) and gentamicin (0.05 mg; s.c.). Rats were allowed to recover for 2-4 wk. Control rats (n=5) experienced the same surgical procedures as experimental rats but without viral infusion.

### Optrode implantation surgery

Two to four weeks after viral infusion surgery, optrodes were implanted into the rNTS, Initially, rats were given buprenorphine-HCl (0.05 mg; s.c.) and atropine sulfate (0.054 mg/kg; s.c.). Animals were then anesthetized with 3% isoflurane in O_2_ at a flowrate of 0.9 L/min and the scalp was shaved. Anesthesia was maintained with 1-3% isoflurane. The rat’s head was placed in a stereotaxic instrument (David Kopf Instruments, Tujunga, CA) and swabbed with betadine and 70% ethanol 3 times. The eyes were lubricated and the rat’s temperature was maintained at 37 °C throughout the surgery. The skull was exposed from just anterior to bregma to about 1.5 cm behind the occipital ridge. Five self-tapping screws were inserted into the skull. The head was angled with bregma 4 mm below lambda and a hole drilled at 14.3-15.3 mm posterior and 1.7-1.8 mm lateral to bregma. The exposed dura was resected and an optrode consisting of 8 or 16 tungsten wires attached to a fiberoptic implant were lowered through the hole to ∼ 5-6 mm below the surface of the brain at a rate of 1mm per 5min. The 16 channel electrode + fiberoptic bundles were drivable and placed ∼ 500µm above the rNTS. A ground wire was wrapped around one of the skull screws. The entire assembly was then embedded in dental acrylic. Rats were administered buprenorphine-HCl (0.05 mg; s.c.) and gentamicin (0.05 mg; s.c.) immediately following surgery and daily for two additional days. The rat was allowed to recover for 5 days or until it regained 90% pre-surgical body weight before testing began.

### Experimental Paradigm

Rats were moderately water deprived (22-23h) and placed in an operant chamber (Med Associates, St. Albans, VT) with free access to a lick spout. Taste stimuli consisted of 0.1 M sucrose, 0.1 M NaCl, 0.1 M MSG:0.01 M IMP, 0.1 M KCl, 0.1 M NH_4_Cl, 0.01 M citric acid, 0.0001M quinine, and artificial saliva (AS; 0.015 M NaCl, 0.022 M KCl, 0.003 M CaCl_2_; 0.0006 M MgCl_2_; pH ∼ 7.4; Hirata et al., 2005; Breza et al., 2010). All tastants were reagent grade and dissolved in AS. Each reinforced lick delivered 12 µL of fluid. Each taste trial consisted of 5 consecutive licks of a taste stimulus separated by 5 licks of an AS rinse presented on a variable ratio 5 schedule. During a randomly-interspersed half of the taste stimulus trials, optogenetic stimulation of GABAergic neurons (473nm; 25 Hz; 10-12 mW) was triggered for 1s after each stimulus lick. The fiberoptic implant was static, but every 2-4 recording days, the microwires were extended ventrally 25-50 µm. Experimental sessions were 30min in length and continued daily, except for weekends, for 2-4 wks.

### Electrophysiological Recording and Light Stimulation

During the experimental session, the rat’s electrode bundle was connected to an Omniplex D Neural Data Acquisition System (Plexon, Dallas, TX). Timing for electrophysiological activity and stimulus events were recorded using PlexControl software (Plexon, Dallas, Tx). The fiberoptic implant was attached to a 473 nm laser source (Shanghai Laser and Optics Century Co., Ltd., Shanghai, China) through a fiber optic patch cable (1m length, 200µm core, 0.22 NA) (THORLABS, Newton, NJ). Optic stimulation was triggered in a random half of tastant trials as mentioned in the experimental paradigm section.

Neuron signals were isolated in Offline Sorter (Plexon, Dallas, TX) or through a semi-supervised spike sorting Python program (Pl2_sort) modified from Mukherjee et al. (2017). The Pl2_sort program contained three scripts of which the user interacted with (Pl2_preprocessing.py, Pl2_postprocessing.py, and Convert_to_nex.py) and a handful of scripts that were called upon; PyPl2 scripts (containing Clustering.py, Pl2_waveforms_datashader.py, pypl2api.py, pypl2lib.py, PL2FileReader.dll, and PL2FileReader64.dll) and a Pl2_processing.py script. The PyPl2 scripts for Python-3 were obtained from Plexon and used to extract data from pl2 files. The Pl2_preprocessing script was used to obtain sorting criteria from the user, select Pl2 files, obtain continuous data and event information from Pl2 files, and save the generated information as an HDF5 file. It then called on the Pl2_processing script using multicore processing and units were isolated from each channel as described in Mukherjee et al. (2017) with slight modifications. The first five principal components were used to determine isolation instead of 3, threshold for significant waveforms was set at 2.5 standard deviations from average voltage value, sampling rate was performed at 40 kH, and 3-7 clusters were generated for the user to select from. After isolation, the Pl2_postprocessing script was used to select isolated units or to further isolate units and then convert the information for waveform shapes, waveform timestamps, events, and event timestamps to json file format. The Convert_to_nex script was used in Neuroexplorer to convert the json file into a Neuroexplorer file for neuron response analysis. Single cells were considered well-isolated if the signal-to noise-ratio was greater than 2, the mean of Mahalanobis distances within a unit waveform cluster was shorter than the mean of Mahalanobis distances between unit waveform clusters, and less than 0.5% of waveforms contained an interspike interval less than 1 ms.

### Analyses of Taste Responses

As in previous work (Escanilla et al., 2015; Roussin et al., 2012; Weiss et al., 2014), responses to taste stimuli were detected over two time scales: 1) responses that extended across more than one lick, called “5-lick” responses, and 2) responses that occurred briefly after each lick, called “lick-by-lick” responses.

5-lick responses were quantified by a significant increase or decrease in firing rate compared to baseline firing rate for at least 300 ms. Baseline firing rate was calculated in 100ms time bins over the 1s preceding the first taste stimulus lick in a trial. To determine if a significant response was present, the firing rate in 100 ms time bins, beginning with the first taste stimulus lick, was compared to the 95% confidence limits of the baseline firing rate. The 100 ms window was moved in 20 ms increments until there were at least three consecutive, non-overlapping 100 ms bins where there was a significant difference between baseline and response firing rates. The leading and trailing edge of the significant bins were used to determine when a taste response started (latency) and ended (duration) respectively. A maximum of two bins within a response were allowed to be non-significant. The response magnitude (firing rate during a response minus the baseline firing rate), latency, duration, and baseline activity were calculated for each taste response. Neurons with a response spike rate less than 2 spikes per sec (sps) were not included.

Lick-by-lick responses were detected using a Chi-squared test comparing responses from the average spike rate of the last non-reinforced lick (i.e. a dry lick) before every tastant trial to the average spike rate of every lick from each tastant. Response windows were limited to 150 ms after each lick and divided into ten 15 ms bins. Using the Chi-squared test, the actual response value from each bin of each tastant vs dry lick was compared to the corresponding expected response bin. A Bonferroni correction was made for multiple (*n* = 16 tastants) comparisons. Neurons with a response firing rate less than 4sps during lick bouts were excluded.

### Analysis of breadth of tuning

In addition to noting the number of tastants to which a cell responds, breadth of tuning was assessed by calculating two standard measures of tuning breadth that are more graded: taste entropy and taste sharpness. Each measure reflects a different aspect of tastant specificity. Both analyses were performed using the five prototypical tastants (sucrose, NaCl, MSG, citric acid, and quinine). Taste entropy (Smith and Travers, 1979) is a measure of uncertainty based on the similarity of response magnitudes between tastants. This measure is calculated as follows:

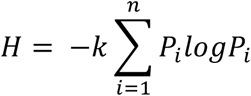

where *n* is the number of tastants (5), *k* = 1.4307 for 5 tastants, and *P_i_* is the ratio of tastant *i* response magnitude to the sum of all tastant response magnitudes. The value ranges from zero, signifying that the neuron responds to a single tastant, to one indicating that the neuron responds to all 5 tastants equally. Taste sharpness (Rainer et al., 1998) is a measure of how similar taste magnitudes are to the best stimulus and is calculated as follows:

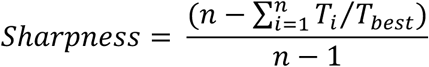

where *n* is the number of tastants, *T_i_* is response magnitude for tastant *i*, and *T_best_* is the response magnitude of the best stimulus. Similar to the entropy measure, a value of zero indicates a response to a single tastant, a value of one indicates equal responses to all five tastants.

### Temporal Coding Analysis

Analysis of information about taste quality conveyed by individual neurons was performed using metric space analyses (MSA; Victor and Purpura 1996, 1997). This method has been described in detail previously (Roussin et al. 2012; Weiss et al. 2014; Escanilla et al. 2015; Sammons et al. 2016) and is only summarized here. The basic approach of MSA is to measure the “cost” of converting one spike train, e.g. a response to a tastant, into another as a measure of similarity/dissimilarity. Cost is accrued by insertion or deletion of spikes or movement of spikes in time. The insertion or deletion of a spike costs one arbitrary unit. Movement of a spike in time costs *qt* units where *q* is a parameter of temporal precision (1/*q* has units of seconds) and *t* is the amount of time that the spike is shifted. Thus, at *q* = 0, the cost of moving a spike is zero, so spike timing is ignored when comparing spike trains; as *q* increases, spike timing is taken into account with progressively greater precision. At each value of *q*, the mutual information *H* between tastants and neural responses is estimated by comparing the similarity of pairs of responses to the same stimulus with the similarity of responses to different stimuli. To mitigate biases due to sample size, the Treves-Panzeri-Miller-Carlton (TPMC) debiaser was applied to all estimates of *H* (for a review see Panzeri et al. 2007). This computation of information conveyed about taste quality is carried out across a range of values of *q,* and the maximum is denoted *H_max_*.

Two auxiliary analyses using synthetic data were also conducted. First, to account for residual bias in the estimation of information, spike trains for 40 pairs of randomly-labeled responses were compared using MSA; this yields *H_shuffled_*. Second, to determine whether temporal information was due to spike timing *per se*, vs. differences in the rate envelope, spikes within each taste-evoked spike trains were randomly assigned to alternative responses to the same tastant, while preserving the rate envelope; calculation of information from these synthetic datasets this yields *H_exchange_*. Information about taste quality conveyed by spike timing was considered significant only if *H_max_* > *H_shuffled_*+2SD and *H_max_* > *H_exchange_*. If *H_max_* > *H_shuffled_*+2SD but not *H_exchange_*, information was considered significant, but information conveyed by spike timing was not considered significant. Information from neurons where *H_max_* ≤ *H_shuffled_*+2SD was set to zero.

To characterize the information conveyed by the population of cells, we calculated the average amount of information conveyed by the entire sample of units at 200, 500, 1000, 1500, and 2000 ms of the cumulative response. Information conveyed by the lick pattern was determined in the same way as for spike trains, and compared with that conveyed by spike trains. Only neurons from sessions that contained at least 6 trials for each tastant were included in the temporal coding analysis.

### Statistical Analyses

*Lick coherence* – For each neuron’s firing pattern, its coherence with the occurrence of licks was calculated using the NeuroExplorer 5.201 Coherence Analysis function (NexTechnologies, Colorado Springs, CO). Single taper Hann windowing was used to calculate the values of 256 frequency bins between 0 and 50 Hz frequency with a 50% overlap between windows. The analysis calculates confidence as described in (Kattla and Lowery, 2010). Neurons with a coherence value above 99% confidence between 4-9 Hz were considered lick coherent. In lick coherent neurons, differences in lick coherence were obtained around tastant licks with optic stimulation versus tastant licks without optic stimulation. The reported difference in coherence value was calculated as the maximum difference in coherence between 4-9 Hz. The reported difference in coherence value was calculated as the maximum difference in coherence between 4-9 Hz. An F-test was used to determine whether the change in coherence observed between baseline conditions without GABA activation and during GABA activation was actually due to GABAergic activation, or random chance. There was no clear distinction between neurons that changed coherence with GABAergic stimulation and those that did not so we chose to split the taste neurons into the three groups based on the value in which 25% of the sample increased or decreased coherence. For the 5 lick neurons (n=40), the 10 that had the greatest decreased in coherence were considered to decrease in coherence and the 10 that had the greatest increase in coherence were considered to increase in coherence. For the lick by lick neurons (*n* = 92), the 23 neurons that had the greatest decrease in coherence were considered to decrease coherence and the 23 neurons that had the greatest increase in coherence were considered to increase coherence. Analyses of the change in coherence for the 5 lick responses and lick by lick responses were independent of each other.

Spearmans’s rank correlation coefficient (*ρ*) was calculated to determine correlations between lick coherence and measures of taste specificity. The 2-tailed *p*-value for each value was obtained for each correlation and a Bonferroni correction was made for multiple (n = 6) comparisons. The six different comparisons were lick coherence versus taste tuning, coherence versus taste entropy, and coherence versus taste sharpness, each with and without optic stimulation.

### Histology/ Immunolabeling

Rats were euthanized with sodium-pentobarbital (390 mg/kg; i.p.). Just before expiration, 10 s of 1 mA DC current was passed through the microwire with the last taste response. The rat was then transcardially perfused with isosaline followed by 4% paraformaldehyde (PFA) in phosphate buffered saline (1x PBS). The brain was extracted and placed in 4% PFA overnight. The next day, brains were washed 3 times with PBS and stored in 20% sucrose in 1x PBS. Brains were then sectioned into 35 µm coronal slices. Every other section was individually placed into wells of a 96 well dish containing a cryoprotectant (30% Ethylene Glycol, 30% Glycerol, 11.4 mM NaH_2_PO_4_-H_2_O, and 38.4 mM Na_2_HPO_4_). The other half of the sections were placed directly onto superfrost plus slides and stained with cresyl violet for lesion site identification. The center of each lesion was taken as the final site of recording.

Sections placed into the cryoprotectant were removed and washed 3 times with 1x PBS. They were placed in blocking agent (10% bovine serum albumin (BSA), 0.1% Triton X, 1x PBS) and gently rocked for 1h at room temperature (RT). Sections were then placed in primary (10% BSA, 1:1,000 Rabbit anti-GFP (Abcam, Cambridge, UK, cat#AB290), 1:500 Mouse anti-NeuN (Millipore, Burlington, MA, cat# MAB377), 1x PBS) for an additional 2h at RT or overnight at 4 °C. Sections were washed 3 times with 1x PBS and placed in secondary (1:500 AF488 conjugated Goat anti-Rabbit (Abcam, Cambridge, UK, Cat# AB150077), 1:500 Cy3 conjugated Donkey anti-mouse (Jackson Immuno Research Labs, West Grove, PA, cat# 715-165-151), 1:10,000 DAPI stain (Millipore, Burlington, MA, cat# 5.08741.0001), 1x PBS) for 1h at RT.

## RESULTS

### General response characteristics

We recorded 113 isolated neurons from the rNTS of freely licking rats with optrode implants. A total of 50 (of 113; 44%) neurons responded to at least one of the eight taste stimuli tested. With GABA stimulation, 43 (of 113; 38%) neurons responded to at least one of the eight taste stimuli. Four neurons were unresponsive without GABA stimulation but showed taste responses with GABA stimulation resulting in a total of 54 neurons that responded to at least one tastant either with or without GABA activation. There were 13 neurons, two of these taste-responsive, that were recorded in non-viral control animals; there was no effect of optical stimulation on these cells. Of the 54 recorded taste neurons, 52 (96%) responded to the five prototypical tastants (sucrose, NaCl, MSG, citric acid, or quinine) while the last two only responded to artificial saliva with GABA stimulation. The average spontaneous firing rate for the population was 18.4 ± 2.8 sps, median = 5.6 sps. The spontaneous firing rate for taste responsive neurons (mean = 16.5 ± 4.3 sps, median = 5.2 sps) was not significantly different from the spontaneous firing rate for non-taste neurons (mean = 20.1 ± 3.7 sps, median = 7.0 sps).

All but one of the taste-responsive neurons were significantly coherent with licking. A total of 97 (86%) of the 113 neurons were lick-coherent. Licking decreased the firing rate for 19 (17%) of the neurons and increased the firing rate for 39 (35%) of the neurons, regardless of whether the licks were reinforced or not.

### GABAergic stimulation changed taste profiles

GABA activation modified taste response magnitudes in 22 rNTS cells. Fig. 1 shows a neuron with 5-lick taste responses to NaCl, KCl, NH_4_Cl, and MSG in the absence of optogenetically-induced GABA activation. GABA stimulation increased the response magnitude for KCl and evoked a significant response to sucrose; there was none without GABA activation. In addition, the response to NH_4_Cl was eliminated and the latency of response for MSG was lengthened. GABA activation had no effect on NaCl, citric acid, quinine, or artificial saliva. Fig. 2 shows a neuron with lick-by-lick taste responses. This neuron was responsive to sucrose, NaCl, citric acid, quinine, MSG and artificial saliva in the absence of GABA activation. GABA stimulation shifted the response latencies for sucrose and artificial saliva and eliminated responses to NaCl, citric acid, and quinine. GABA stimulation also generated a response to KCl and had no effect on responses to MSG or NH_4_Cl.

**Fig. 1.**
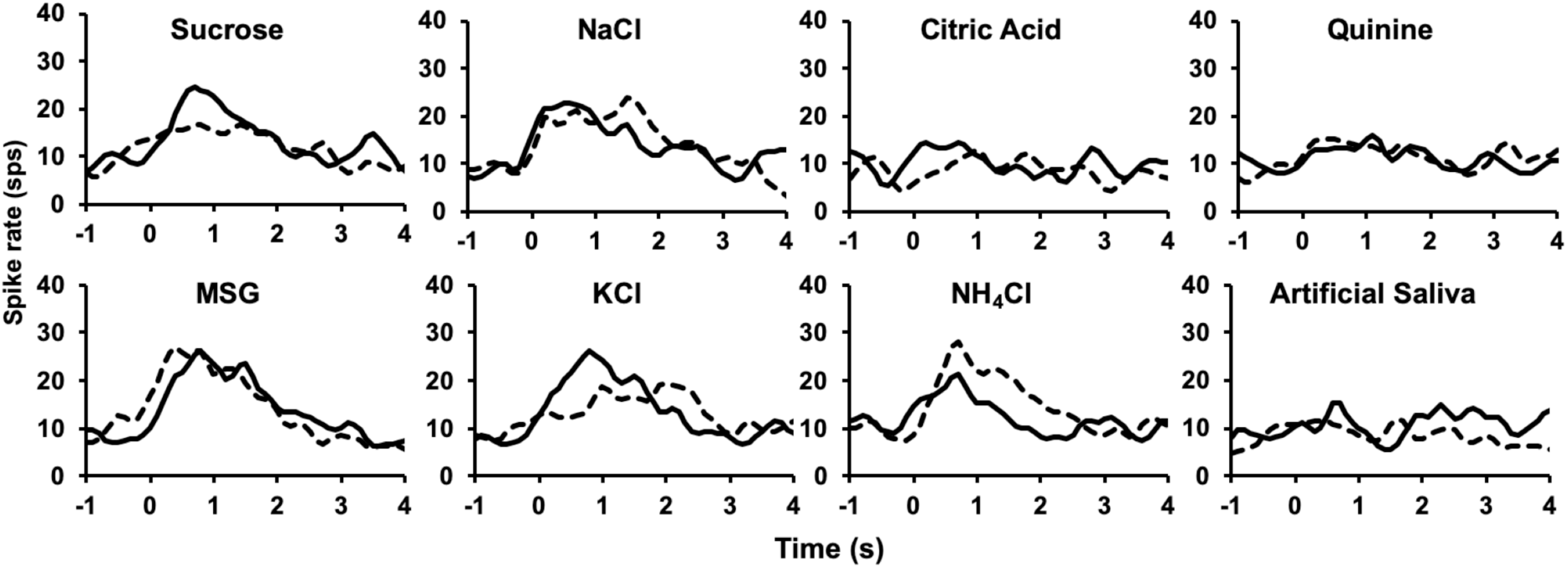
Taste responsive neuron responding on a 5-lick time scale. Histograms show the responses to eight taste stimuli without GABA stimulation (dashed line) and with GABA stimulation (solid line). Without GABA activation, this neuron responded to NaCl, KCl, NH_4_Cl, and MSG. With GABA activation, this neuron changed its taste response profile to include sucrose. The KCl response was enhanced, the NH_4_Cl response was attenuated and the latency for MSG was decreased upon GABAergic stimulation. Responses to citric acid, quinine and artificial saliva were unaffected by GABA stimulation.

**Fig. 2.**
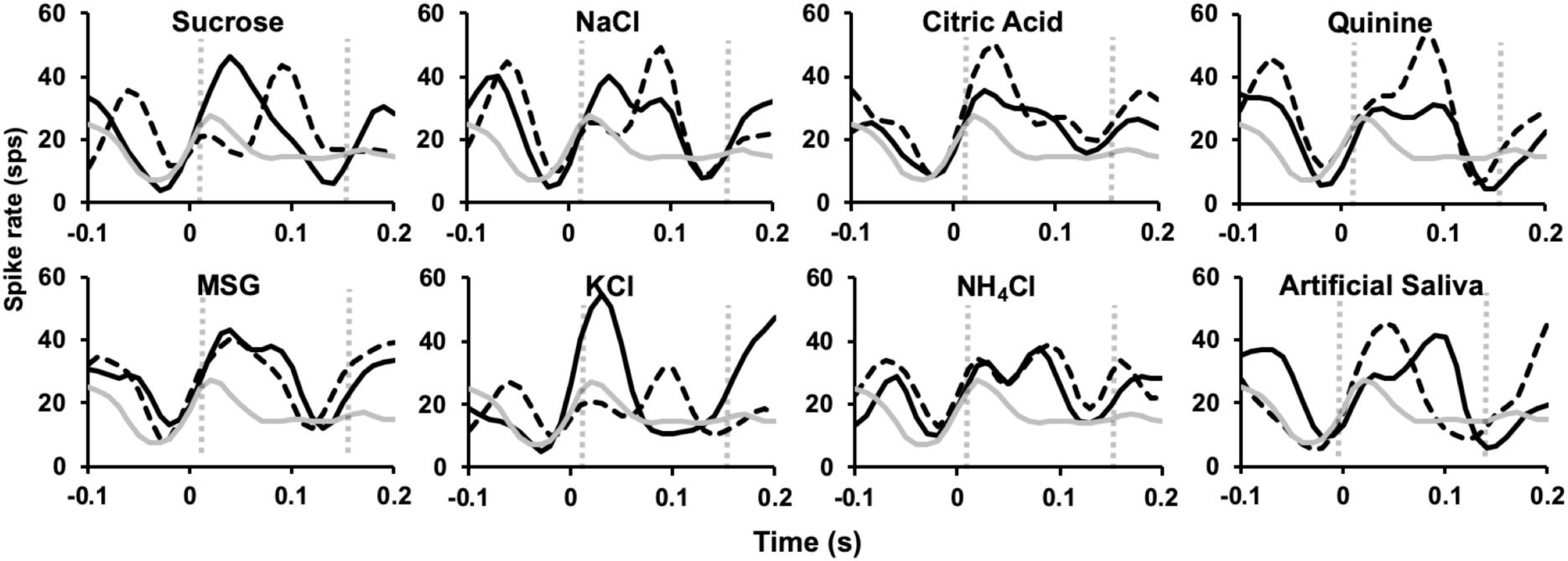
Taste responsive neuron responding on a lick-by-lick time scale. Histograms show lick-by-lick responses to eight taste stimuli without GABA stimulation (dashed line) and with GABA stimulation (solid line); grey line shows response to dry (unreinforced) licks. Without GABA activation, this neuron responded to sucrose, NaCl, MSG, citric acid, quinine, and artificial saliva. With GABAergic stimulation, the latencies of responses were shortened for sucrose and NaCl and lengthened for artificial saliva; responses to citric acid and quinine were attenuated, and a response to KCl appeared. GABA activation had no effect on responses to MSG and NH_4_Cl.

Stimulation of GABA release in the rNTS changed taste response magnitudes for both 5-Lick (Fig. 3A) and lick-by-lick responses (Fig. 3B). Neurons were organized according to their best stimulus, i.e. the stimulus that evoked the largest response without GABA stimulation (grey bars). Responses to tastants with GABA stimulation are overlaid as diamonds. Twenty-two of the 54 (47%) taste taste-responsive neurons had responses that were significantly affected by GABA stimulation (Table 1).

**Fig. 3.**
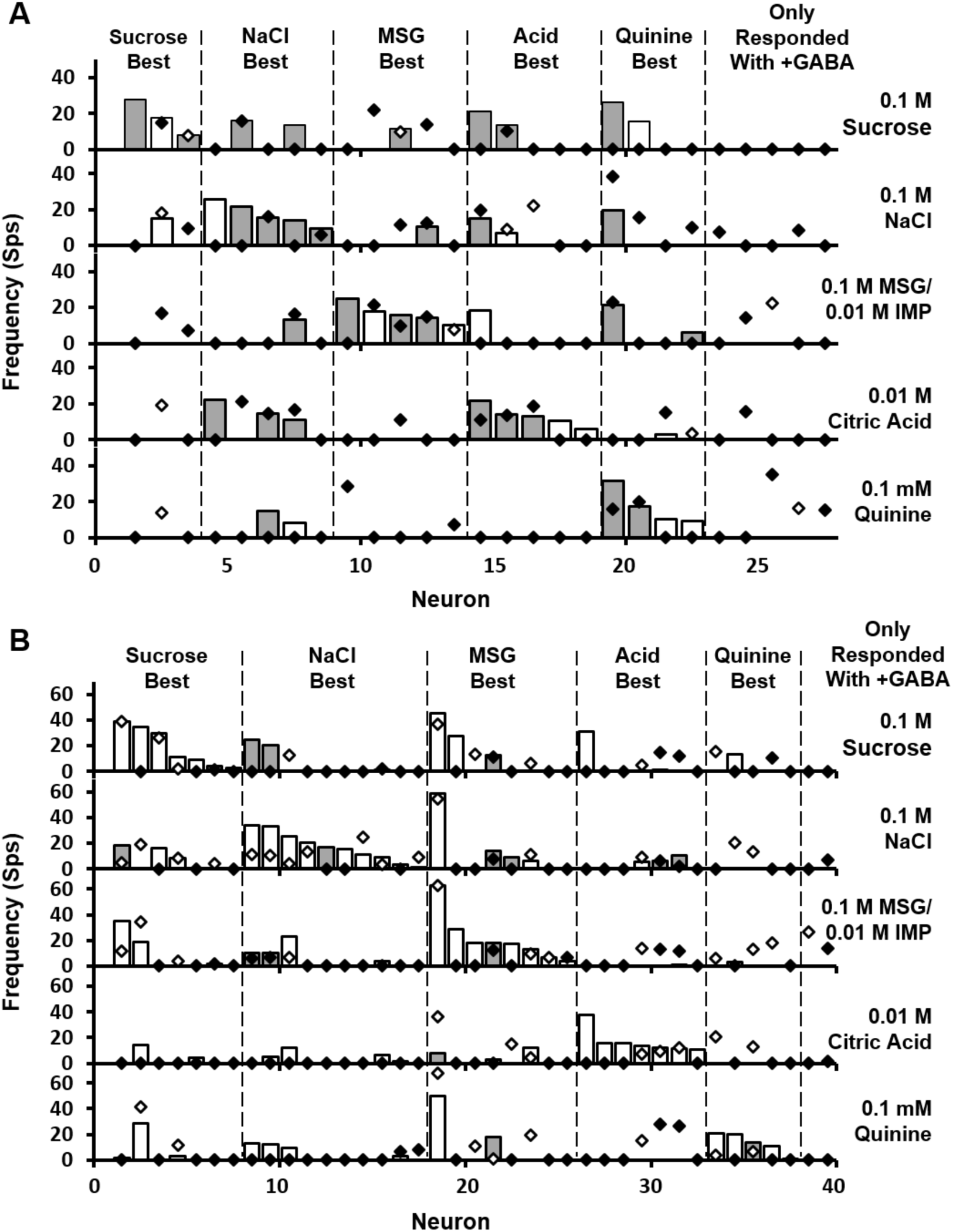
Taste responses of rNTS neurons with and without GABA stimulation. The absolute value of each response was calculated for the 27 5-lick and 41 lick-by-lick taste neurons. Filled bars/diamonds indicate excitatory responses; empty bars/diamonds indicate inhibitory responses. Thirteen neurons responded on both time scales. Neurons were separated by their best-stimulus response without GABA stimulation (grey bars). Responses during GABA stimulation (diamonds) are overlaid on the non-stimulated responses. Neurons that showed taste responses only during GABA stimulation are shown at the far right.

**Table 1.**
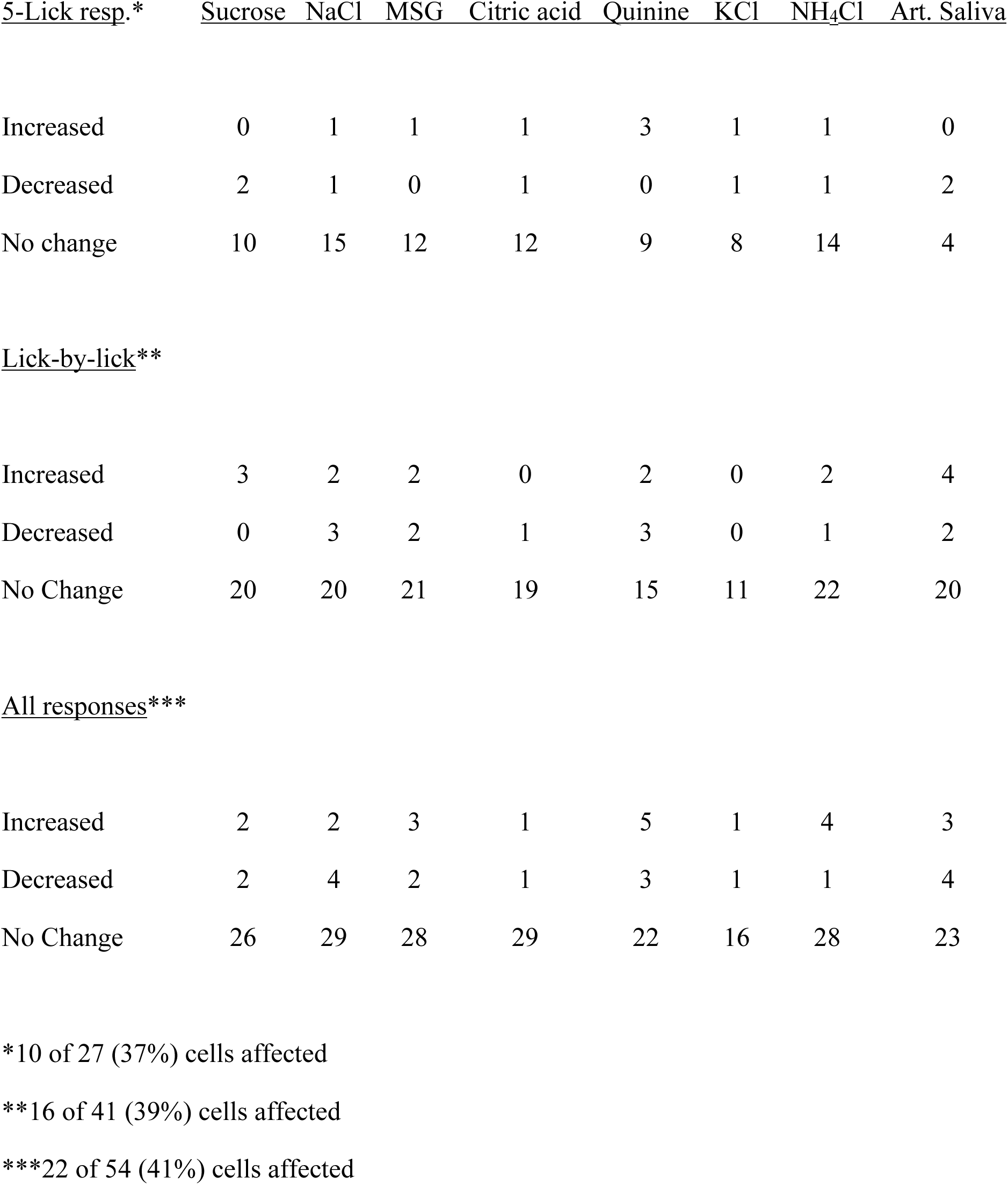
Effect of GABA stimulation on taste response magnitudes in rNTS.

| <u>5-Lick resp.*</u> | <u>Sucrose</u> | <u>NaCl</u> | <u>MSG</u> | <u>Citric acid</u> | <u>Quinine</u> | <u>KCl</u> | <u>NH<sub>4</sub>Cl</u> | <u>Art. Saliva</u> |
| --- | --- | --- | --- | --- | --- | --- | --- | --- |
| Increased | 0 | 1 | 1 | 1 | 3 | 1 | 1 | 0 |
| Decreased | 2 | 1 | 0 | 1 | 0 | 1 | 1 | 2 |
| No change | 10 | 15 | 12 | 12 | 9 | 8 | 14 | 4 |
| Lick-by-lick\*\* |  |  |  |  |  |  |  |  |
| Increased | 3 | 2 | 2 | 0 | 2 | 0 | 2 | 4 |
| Decreased | 0 | 3 | 2 | 1 | 3 | 0 | 1 | 2 |
| No Change | 20 | 20 | 21 | 19 | 15 | 11 | 22 | 20 |
| All responses\*\*\* |  |  |  |  |  |  |  |  |
| Increased | 2 | 2 | 3 | 1 | 5 | 1 | 4 | 3 |
| Decreased | 2 | 4 | 2 | 1 | 3 | 1 | 1 | 4 |
| No Change | 26 | 29 | 28 | 29 | 22 | 16 | 28 | 23 |

\*10 of 27 (37%) cells affected
\*\*16 of 41 (39%) cells affected
\*\*\*22 of 54 (41%) cells affected

Analyses of breadth of tuning across the tastants representing the five basic taste qualities suggested that GABA activation reduced the number of tastants to which a neuron responded. Fig. 4 shows that, although the total number of responses to any given tastant was not altered by GABA stimulation (Chi-square = 1.46, *df* = 4, *p* = 0.835), the number of tastants to which individual neurons responded decreased significantly (Chi-square = 13.62, *df* = 5, *p* = 0.018). In contrast, there were no significant differences in the Uncertainty measure with or without GABA activation. The average taste entropy was 0.37 ± 0.05 without GABA stimulation and 0.48 ± 0.05 (Student’s *t* test, *p* = 0.535) with GABA stimulation. Moreover, GABA activation did not change taste sharpness significantly. Average taste sharpness was 0.81 ± 0.03 without GABA stimulation and 0.76 ± 0.03 with GABA stimulation (Student’s *t* test, *p* = 0.698). add explanation

**Fig. 4.**
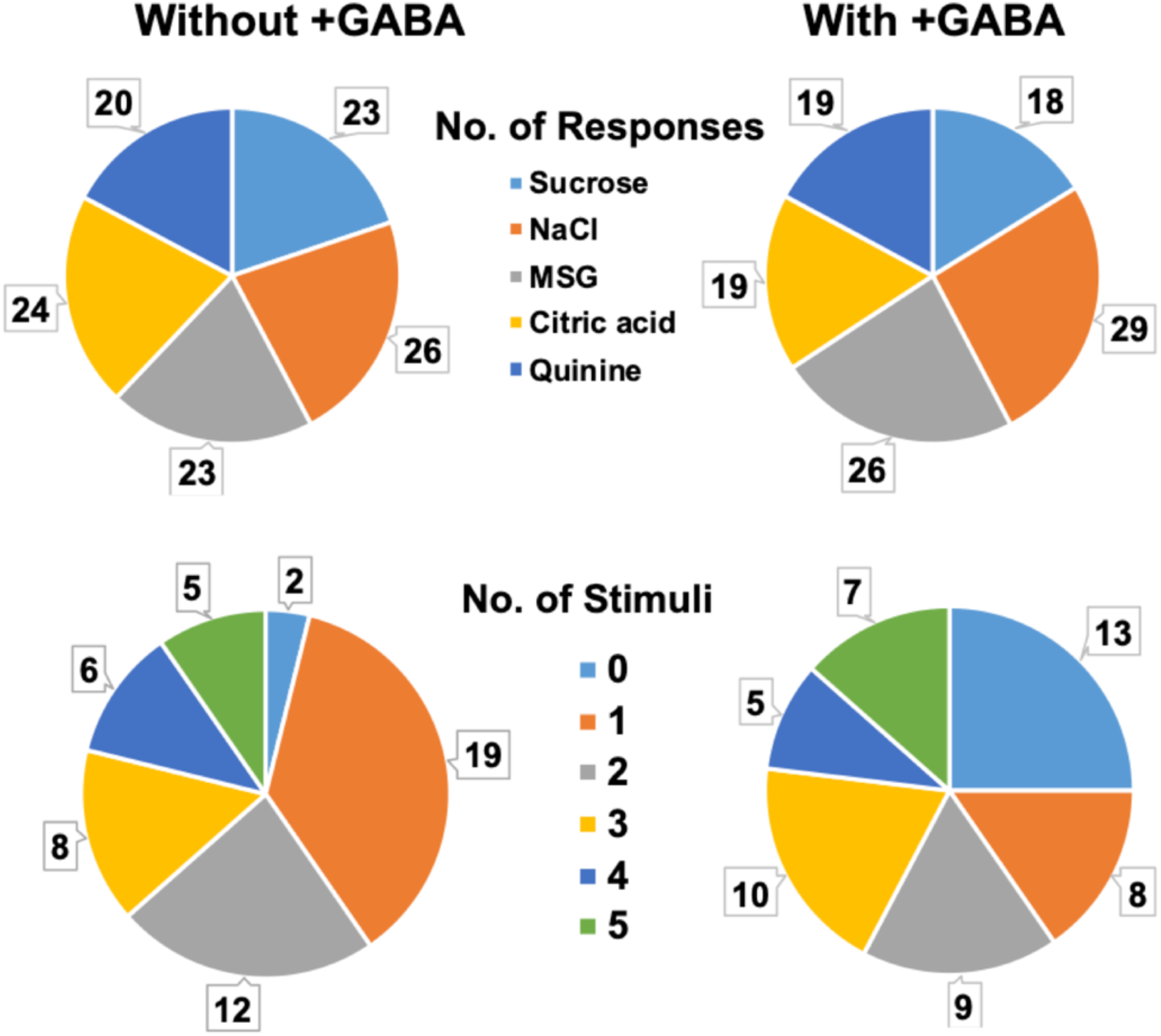
Pie graphs showing the distribution of the number of responses to each of the five basic tastants (top) and the number of tastants to which a neuron responds (bottom). Graphs on the left are distributions without GAB activation; graphs on the right show the distribution s with GABA activation. Although each tastant evoked about the same number of responses across the sample, neurons were significantly more narrowly tuned with GABA stimulation. See text for details.

To assess the relationship among across unit patterns of taste responses, we applied a multidimensional scaling analysis using Pearson correlations as measures of similarity. A hypothetical “taste space” placed the across unit patterns for each taste stimulus close together or far apart depending on their similarity/dissimilarity. Across unit response patterns both before and during GABA activation were analyzed and graphed together. Fig. 5 shows the results of those analyses. Without GABA activation, response patterns to each of the five basic were well separated in taste space suggesting that each tastant evoked easily discriminable patterns of response. With GABA activation, the configuration of across unit patterns was similar but shifted in space, indicating that the basic interrelationships among tastant-evoked response patterns was intact, but the identities of the units that contributed most to the pattern were different.

**Fig. 5.**
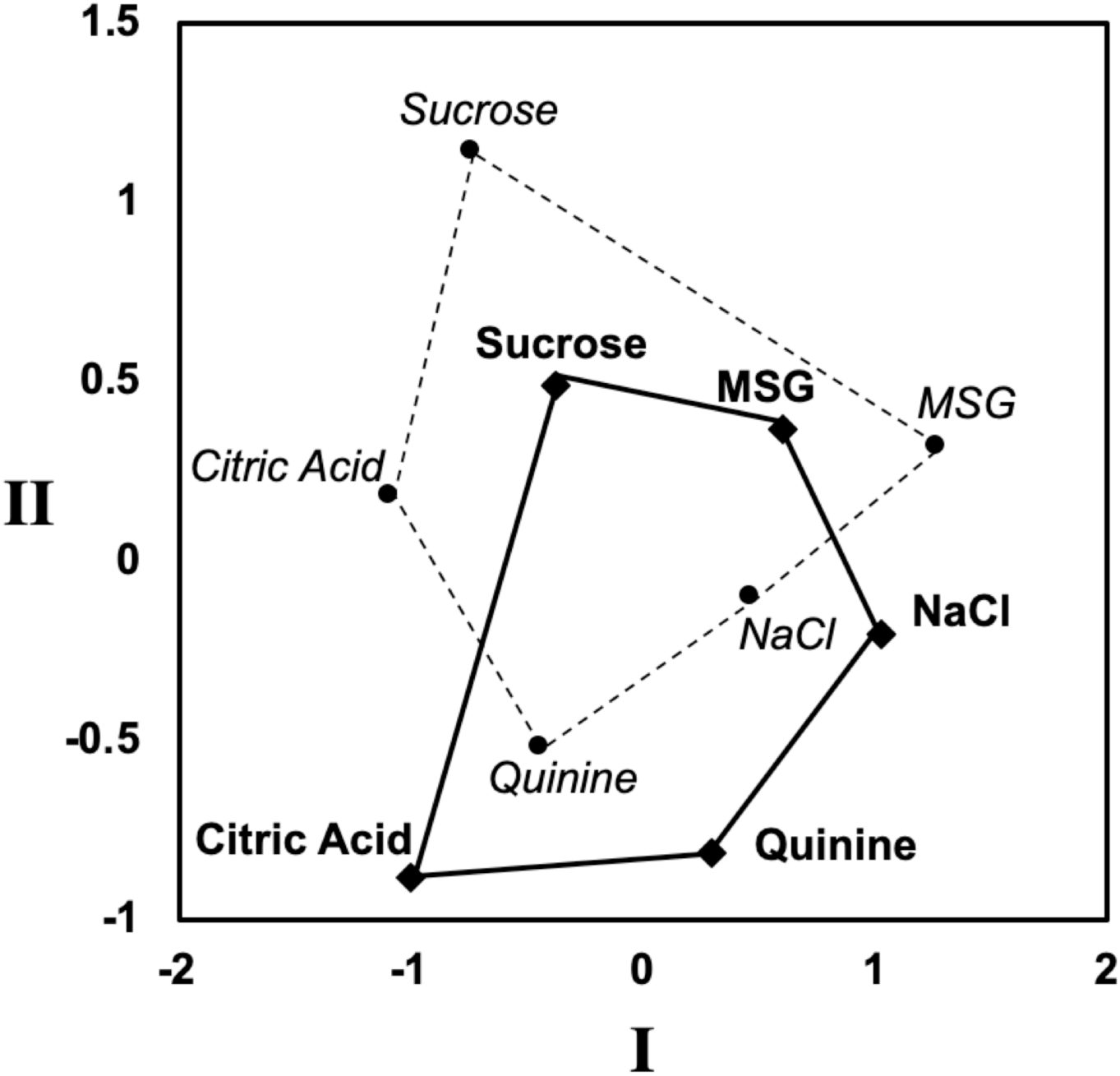
Results of Multidimensional scaling analysis of taste response magnitudes. Pearson’s correlations were used as a measure of similarity for across unit patterns evoked by tastants. A dashed line connects the patterns evoked by taste stimuli without GABA stimulation; a solid line connects the patterns evoked by taste stimuli with GABA stimulation. Gutman stress values were: 1 dimension, 0.281; 2 dimensions, 0.108; 3 dimensions, 0.062; 4 dimensions, 0.026; 5 dimensions, 0.017. GABA activation shifted the location of all taste-evoked across unit patterns, but the overall organization was unchanged.

### Taste-responsive neurons have higher lick coherence than non-taste neurons

The majority of rNTS neurons (97 of 113; 86%) were coherent with licking. Coherence values associated with all licking within a session will be termed overall lick coherence. In general, overall lick coherence values for taste neurons were significantly higher than those of non-taste neurons (*p* < 0.001; taste neurons: mean = 2.2*10^-1^, median=1.7*10^-1^, *n* = 54; non-taste neurons: mean = 6.3*10^-2^, median=2.4*10^-2^, *n* = 59). To determine whether and how GABA activation affected lick coherence, we restricted coherence analysis to licks that resulted in taste stimulus delivery, since this is when GABA release was triggered. Coherence values associated with licking only during tastant delivery will be termed taste-restricted lick coherence Not surprisingly, tastant-restricted lick coherence values without GABAergic stimulation between taste neurons and non-taste neurons were also significantly different (*p* <0.001; taste neurons: mean = 3.3*10^-2^, median = 2.3*10^-2^; non-taste neurons: mean = 1.0*10^-2^, median=5.2*10^-3^). The distribution of lick coherence values for both the global lick coherence and tastant-restricted lick coherence can be seen on the x-axes of Figs. 6A and 6B, respectively. Figure 6B additionally tracks the change in tastant-restricted lick coherence upon GABAergic stimulation over the y-axis and shows that taste responsive neurons are affected to a greater degree than non-taste neurons. A two sample F-test on the amount that GABA stimulation changes tastant-restricted lick coherence in taste neurons vs. non-taste neurons shows a significant difference in the variance of the GABA effect (*p* < 0.001).

**Fig. 6.**
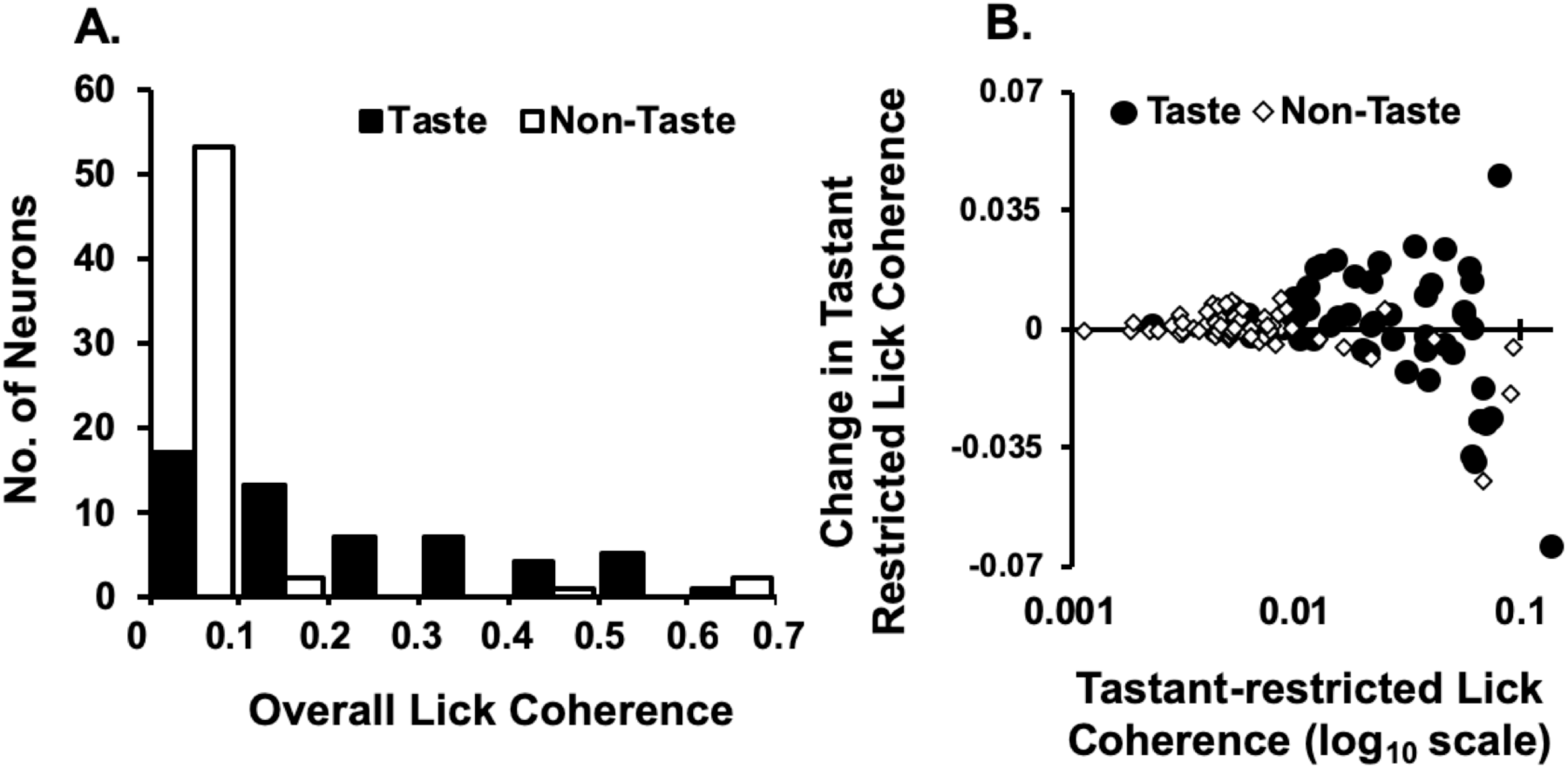
Lick coherence in taste-responsive and non-taste-responsive neurons. **A.** Distribution of overall lick coherence values for both taste (filled bars) and non-taste (hollow bars) neurons. Overall lick coherence values were calculated using all licks whether the lick was reinforced or not. **B.** Change in tastant-restricted lick coherence values with GABA stimulation (y-axis) with taste-restricted lick coherence without GABA activation shown on the abscissa.

### GABA activation increased gustatory information in rNTS neurons

To analyze the effect of enhancing GABAergic tone on temporal coding of taste stimuli, we applied MSA to datasets with at least 6 repetitions of each tastant (with and without GABA activation). Sixty neurons (38 taste-responsive; 22 not taste responsive) were included in these analyses; as previously noted (Denman et al., 2019), neurons that are not considered “taste-responsive” by classical criteria nevertheless may carry information about taste when analyzed by MSA. Among the 38 taste-responsive neurons, GABA stimulation reduced taste-related information to zero in 10 and generated significant taste-related information in 14 (Fig. 7A). Previous work has shown that spike patterns in neurons that do not generate enough taste-evoked spikes to be considered taste-responsive may nevertheless convey information about taste quality in the temporal arrangement of spikes following a taste stimulus (Denman et al., 2019). GABA activation eliminated taste-related information from nine of the 22 non-taste-responsive neurons and generated information from five non-taste-responsive neurons (Fig. 7B).

**Fig. 7.**
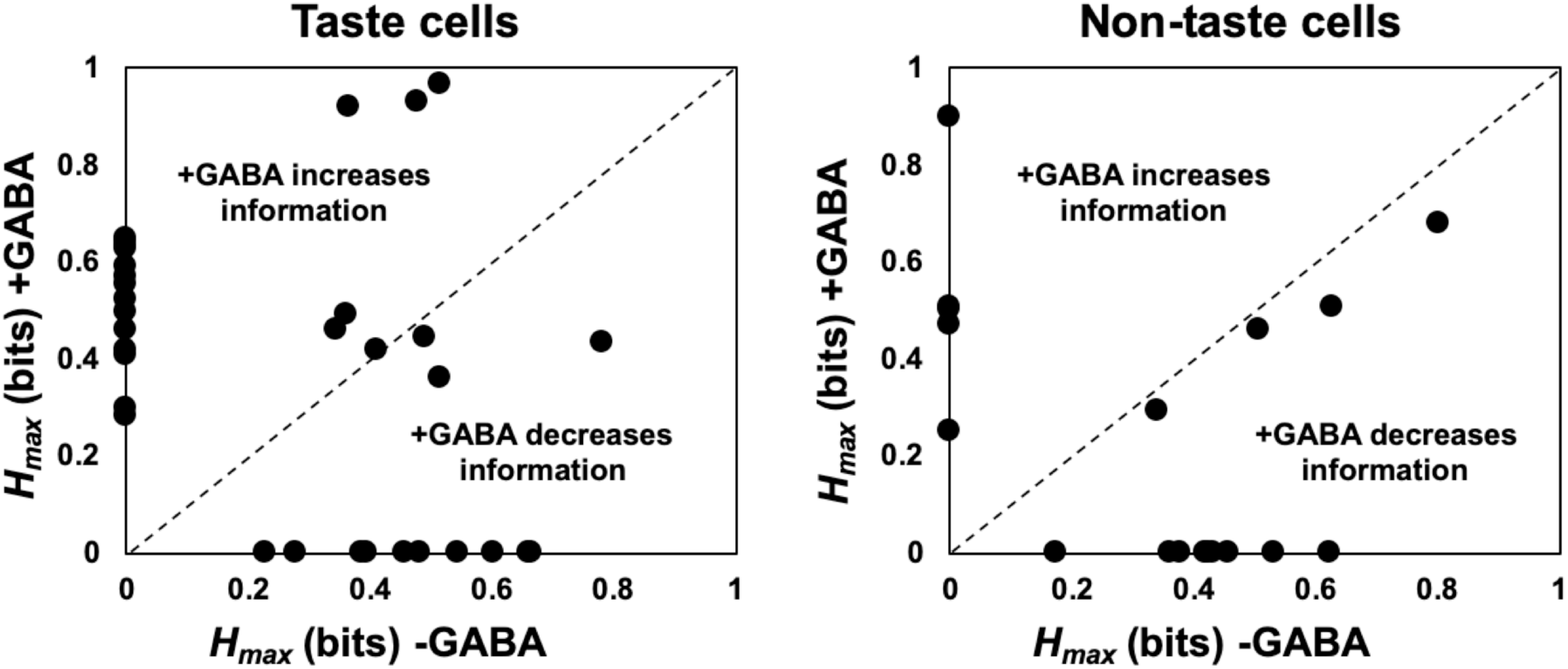
Information (in bits) conveyed about taste quality in taste and non-taste neurons in rNTS. For each cell, *H_max_* was plotted without vs. with laser-stimulated GABA release. **A.** taste cells or **B**. non-taste cells. Only neurons with at least 6 trials for each tastant were used. Information was conveyed by the temporal aspects of taste responses in 33 of 54 (61%) taste cells and 18 of 59 (31%) of non-taste cells. GABA stimulation either enhanced or attenuated information conveyed about taste quality in 73% (24 of 33) taste cells and 78% (4 of 18) non-taste cells.

Taste-related information conveyed by spike timing was also analyzed at various response intervals ranging between 200ms to 2s. Fig. 8 shows the results of those analyses. At 2s, GABA activation increased taste-related information on average by 0.11 bits (48% increase) when all cells with 5-lick taste responses were considered. Because taste-restricted lick coherence was modified in some cells by GABA stimulation, we divided all the neurons (*n* = 113) into quartiles based on the change in GABA-evoked changes in taste-restricted lick coherence. Cells in the uppermost quartile in which GABAergic stimulation increased lick coherence (*n* = 28) had a consistent increase in taste information with a maximum increase of 0.16 bits (68% increase) at 2s with GABA activation. Taste-related information was not affected by GABA stimulation in neurons in the bottom three quartiles. Further, information on the rats’ lick patterns of different taste qualities was overall slightly decreased by GABA activation, suggesting that changes in the lick pattern *per se* cannot account for the increased taste quality information.

**Fig. 8.**
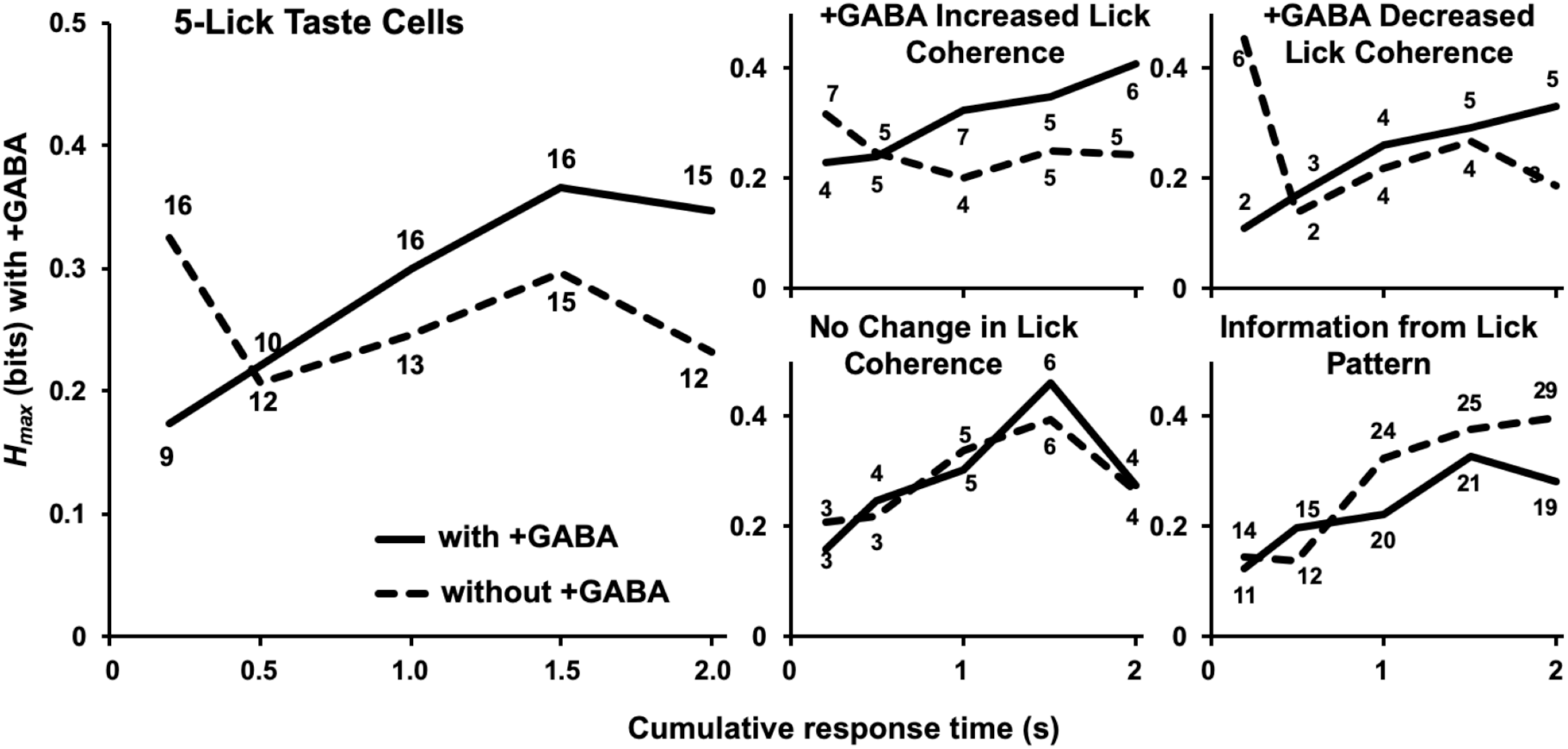
Information (in bits)conveyed about the five prototypical tastants from the population of rNTS neurons over the first 2s of response. Separate analyses were conducted at each response interval. Temporal coding information obtained with (solid line) and without (dashed line) GABA stimulation is shown. **Left.** Information for all taste ells with 5-lick responses. **Right.** Neurons were separated into groups depending on how GABA stimulation affected stimulus-restricted lick coherence. Also shown is the effect of GABA stimulation on the taste-related information conveyed by the lick pattern. Numbers adjacent to each data point denote number of neurons in which significant taste quality information was obtained. GABA stimulation enhanced the information conveyed about salty tastes only in those cells where GABA also enhanced lick coherence. GABA stimulation did not affect the information conveyed by the lick pattern.

### Information about salty tastants is increased after GABAergic stimulation

In addition to the five prototypical tastants, we also tested KCl and NH_4_Cl to determine if GABA stimulation would increase the distinction between tastants of the same taste quality. Over the entire population of neurons, GABAergic stimulation had no effect on information relayed on the salty tastants (Fig. 9). However, when broken into the effect of GABAergic stimulation on lick coherence, similar to information about taste qualities, information relayed about the salty tastants was increased when lick coherence was also increased (Fig. 9) with a maximum increase of 0.13 bits (51%) at 1.5 s. Similar to taste quality analysis, GABAergic stimulation had no effect on neurons with decreased coherence or no change in coherence and it had no effect on information from the lick pattern.

**Fig. 9.**
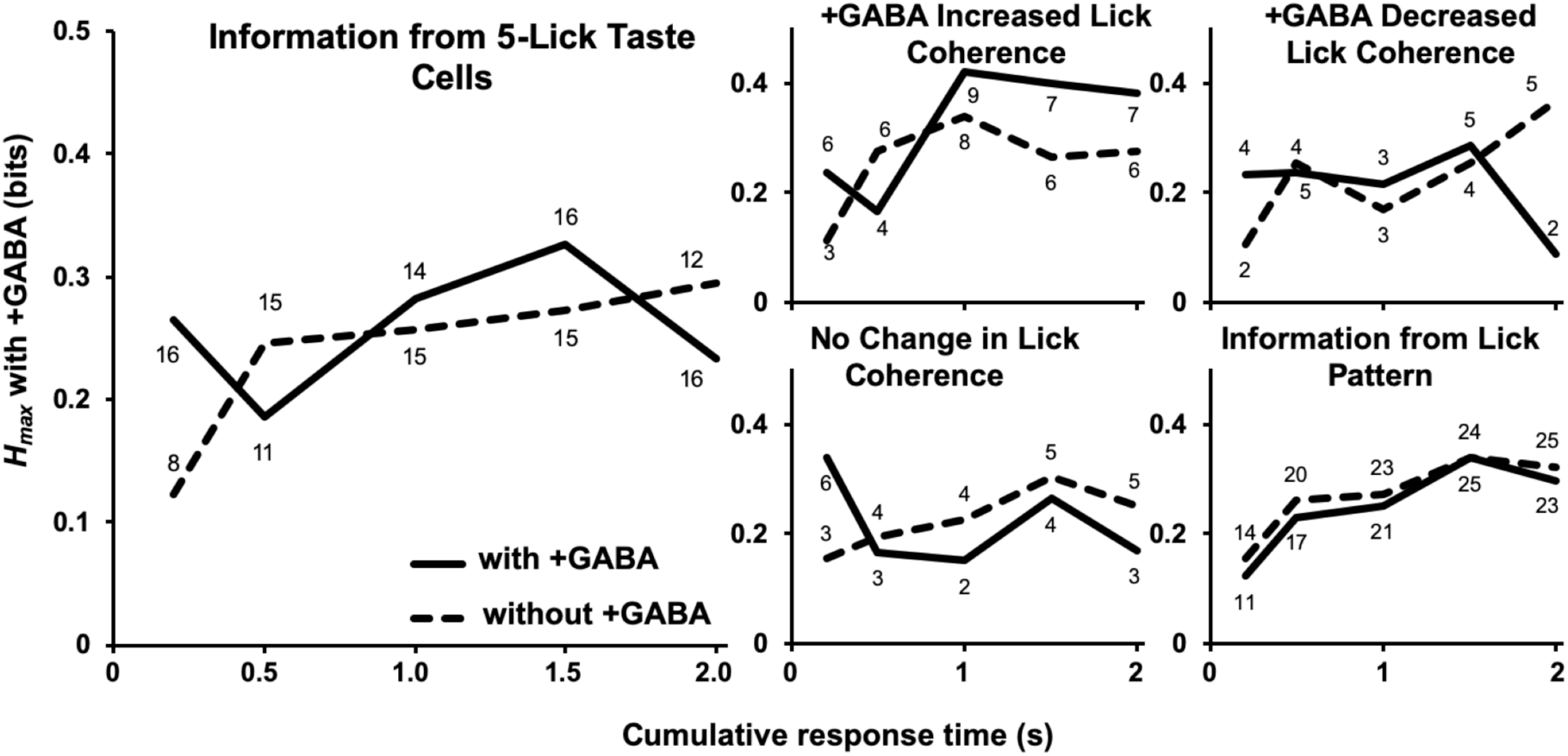
Information (in bits) conveyed about salty tastants (NaCl, KCl, and NH_4_Cl) from the population of rNTS neurons over the first 2s of response. Separate analyses were conducted at each response interval. Temporal coding information obtained with (solid line) and without (dashed line) GABA stimulation is shown. **Left.** Information for all taste ells with 5-lick responses. **Right.** Neurons were separated into groups depending on how GABA stimulation affected stimulus-restricted lick coherence. Also shown is the effect of GABA stimulation on the taste-related information conveyed by the lick pattern. Numbers adjacent to each data point denote number of neurons in which significant taste quality information was obtained. GABA stimulation enhanced the information conveyed about salty tastes only in those cells where GABA also enhanced lick coherence. GABA stimulation did not affect the information conveyed by the lick pattern.

### Effect of GABA activation on information about palatability

Much of the information increase obtained by GABA stimulation occurs a second or two after tastant delivery is initiated. This time epoch is thought to signal taste palatability, at least in the gustatory cortex (Katz et al. 2001). As such, we sought to determine if GABAergic stimulation would increase information about the palatability of tastants. We collapsed responses to sucrose and NaCl as the palatable tastants and collapsed responses to citric acid and quinine as the non-palatable tastants and performed MSA on the two groups. Once again, GABAergic stimulation increased information in the later time points (Fig. 10) with a maximum increase of 0.21 bits (118%) at 1.5 s. GABA-induced increase in palatability-related information was observed whether GABA activation increased lick coherence (maximum 0.27 (246%) at 1.5s), decreased lick coherence (0.11 (58%) at 1.5s) or had no effect on lick coherence (0.23 (86%) at 1.5s), with a non-significant trend to greater increases in palatability-related information in those neurons in which GABA had a larger effect on lick coherence. Again, information conveyed solely by the lick pattern was not changed with GABAergic stimulation.

**Fig. 10.**
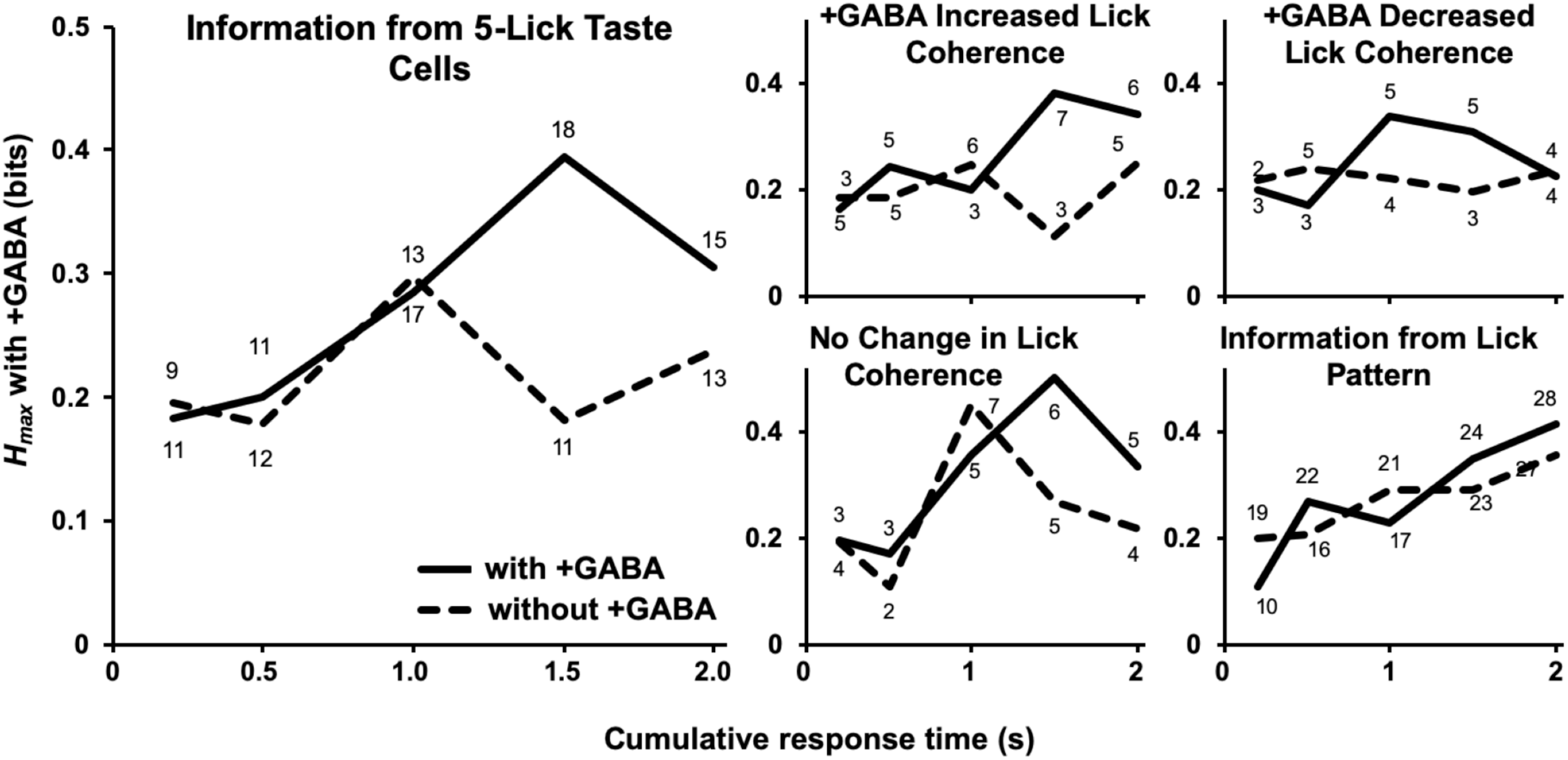
Information (in bits) conveyed about palatable (sucrose, NaCl) vs. unpalatable (citric acid, quinine) from the population of rNTS neurons over the first 2s of response. Separate analyses were conducted at each response interval. Temporal coding information obtained with (solid line) and without (dashed line) GABA stimulation is shown. **Left.** Information for all taste ells with 5-lick responses. **Right.** Neurons were separated into groups depending on how GABA stimulation affected stimulus-restricted lick coherence. Also shown is the effect of GABA stimulation on the taste-related information conveyed by the lick pattern. Numbers adjacent to each data point denote number of neurons in which significant taste quality information was obtained. Regardless of the effect of GABA stimulation on lick coherence, GABA stimulation enhanced the information conveyed about taste palatability in taste responses >1s. GABA stimulation did not affect the information conveyed by the lick pattern.

### Histology

Lesion analysis show that the electrodes were dispersed throughout the rNTS from 11.76 to 12.48. The lesions were mostly lateral with the most rostral lesion also being the most medial (Fig. 11A). Figure 11B shows channelrhodopsin expression in the area surrounding the rNTS lesion. Three experimental rats were found to have their fiberoptic implant >1 mm lateral to the NTS and were excluded from the dataset.

**Fig. 11.**
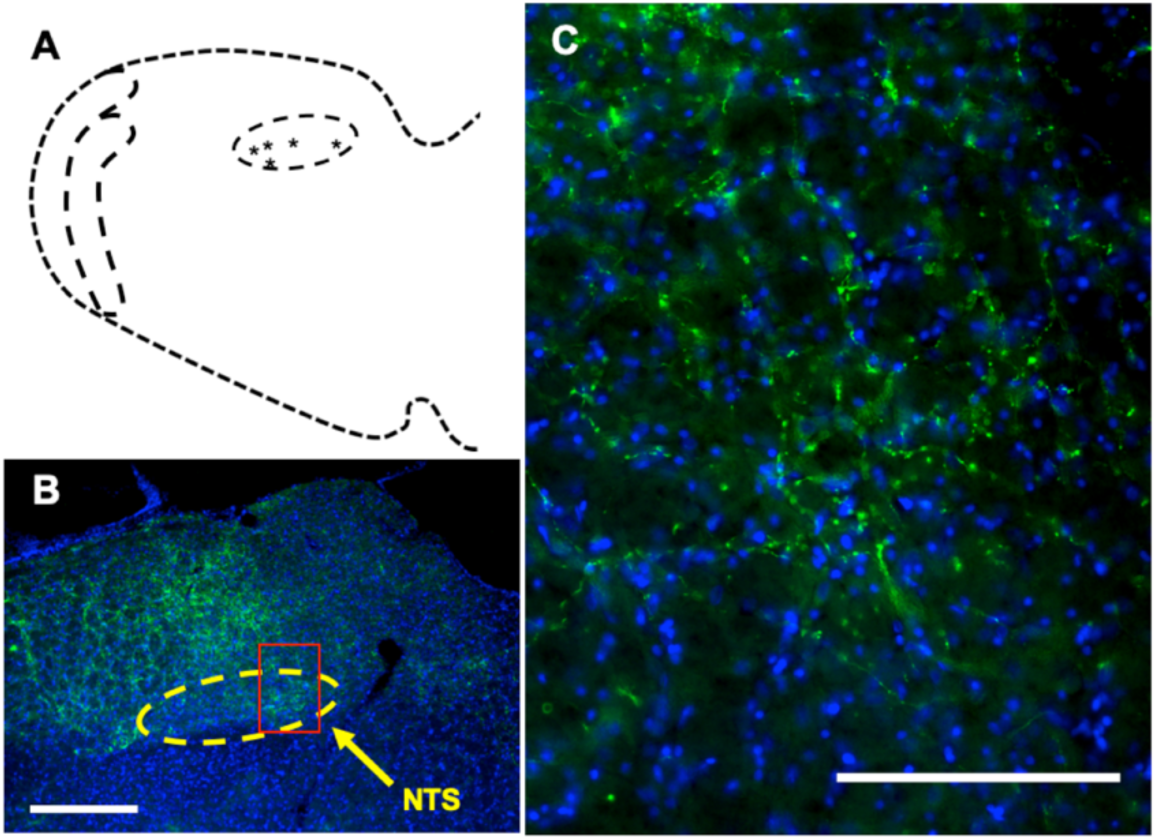
Histological reconstruction of neuronal recordings and channelrhodopsin expression in the rNTS. **A**. Schematic diagram of the brainstem with a dashed oval outlining the rNTS and the center of each lesion represented by an *. Lesions ranged from 11.76-12.48 mm posterior to bregma. **B.** Image of the rNTS (dashed yellow oval); red box represents magnified inset. White scale bar represents 500µm. **C.** Magnified image of rNTS. Channelrhodopsin: green; DAPI: blue. White scale bar represents 100µm.

## DISCUSSION

Enhanced GABAergic tone in the rNTS remodeled the across-neuron pattern of taste responsiveness and enhanced the information discriminating palatable vs. unpalatable tastants conveyed by the temporal characteristics of the response. In individual cells in the rNTS taste response profiles were changed by optical stimulation of GABA terminals in about half (22 of 54; 47%) of the sample of neurons. Responses to some stimuli were enhanced and others attenuated, sometimes within the same cell. In fact, there were neurons that responded to taste stimuli only during GABAergic stimulation (*n* = 2) and others that were rendered completely unresponsive to taste under GABAergic influence (*n* = 13). These GABA-induced cell-by-cell changes did not shift the overall interrelationship among response patterns but instead changed the identities of the cells that contributed to the across unit patterns of response associated with each taste stimulus. Enhancing GABA did, however, change the temporal patterns of taste-evoked activity such that the information discriminating palatable (sucrose and NaCl) vs. unpalatable (citric acid and quinine) increased when longer (1-2s) taste responses were considered.

Present results expand the results reported by Smith and Li (1998). In that study, either GABA or the GABA antagonist bicuculline methiodide (BICM) BICM was infused directly into the rNTS in urethane-anesthetized hamsters. Their data inferred that GABA narrows the tuning of taste-responsive cells in the rNTS. In awake rats in the present study, we confirmed that optical activation of GABA in rNTS narrows taste tuning in a subset of cells; however, it also broadened the response profile in another subset of cells. While Smith and Li (1998) only tested the best and second best stimulus with GABA and BICM, we tested the effects of GABA enhancement for all of the basic tastants and found more complex effects. So, for example, it was not uncommon for GABA enhancement to attenuate the response to one stimulus and amplify the response to another in the same cell. In those cases, the breadth of tuning did not show a net change, though the complement of tastants that evoked a response was altered.

Results of the MDS analyses illustrate the effect of augmenting GABAergic tone on the population coding of taste in NTS. Specifically, under the influence of enhanced GABA release, the taste space generated by the pattern of responsiveness across units was essentially unchanged compared to the taste space without GABA enhancement. That is, in both taste spaces, patterns associated with the five basic taste qualities were well separated from each other; however, the placement of taste stimuli in the taste space with GABA enhancement was systematically shifted. This result implies that, for any given tastant, the identity of the cells that conveyed the signal was shifted by GABA enhancement but the overall signal across tastants conveyed by the population was essentially intact.

While GABA enhancement affected taste response magnitudes, i.e. spike count, it also modified the temporal arrangement of spikes within taste responses. Furthermore, these effects were correlated with GABA-induced changes in lick coherence. In general, GABA enhancement boosted the information conveyed about the five basic taste qualities. A closer analysis suggested that this effect was most prominent in those cells that showed an increase in GABA-induced lick coherence. Moreover, the information conveyed about the three salts that were tested, NaCl, KCl and NH_4_Cl, was increased by GABA enhancement only in those cells where GABA also increased lick coherence. By far, the largest effect of increasing GABAergic tone was seen in the discrimination of palatable (sucrose and NaCl) vs. unpalatable (citric acid and quinine) tastants with the temporal arrangement of tastant-evoked spikes. This effect was apparent regardless of the effect of GABA on lick coherence, with a trend to being larger when this effect was present, and only manifested as longer taste response intervals were considered. Interestingly, it is during the longer response intervals when taste palatability is encoded in the gustatory cortex (Katz et al. 2001); our finding of a similar time-dependence in the brainstem suggests that the action of GABA in the brainstem may be involved in this effect.

GABA enhancement in NTS was found to alter lick coherence during taste stimulus presentation only in taste-responsive cells. This type of coherence is common in the brainstem taste areas of awake unrestrained animals (Denman et al. 2019; Roussin et al. 2012; Weiss et al. 2014); most NTS cells, including most taste-responsive cells, show some degree of lick coherence. We have found that, in addition to taste-responsive cells, non-taste-responsive cells that show significant lick coherence can also convey some information about taste quality (Denman et al., 2019; Roussin et al., 2012; Weiss et al., 2014). Thus, the lick pattern, as reflected in the lick coherent spiking of these cells, can buttress the information about taste quality conveyed by taste-evoked activity. Our data suggest that GABAergic activity may modulate taste-related lick coherence to amplify the contributions of some cells while diminishing the contributions of others to the neural representation of taste in the rNTS, essentially reconfiguring the sensorimotor balance among taste-responsive neurons in rNTS.

The effects of GABA activation reported here must be considered in the context of some obvious limitations. For example, the amplification of GABAergic tone vis optogenetic stimulation is a non-physiological manipulation. Under normal physiological conditions, cells in the rNTS are under a tonic inhibitory influence, with GABA as a major contributor (Grabauskas and Bradley, 2003; Liu et al., 1993; Smith and Li, 1998). Moreover, taste simulation may evoke GABA release in rNTS. Experimental augmentation of GABA release during taste stimulation represents at best a crude exaggeration of the natural influence of GABA on rNTS cells. Nevertheless, the fact that there were systematic effects on taste responsivity both on an individual cell and on a population level implies that there are meaningful concepts that can be derived from our results.

The fact that global optical activation of GABA activates GABA release from a variety of sources represents another limitation of the present study. GABAergic projections arise from both local interneurons in NTS (Davis 1993; Lasiter and Kachele, 1988) as well as centrifugal structures such as the gustatory cortex (GC; Smith and Li, 2000; Torrealba and Muller, 1996) or amygdala (AMG; Batten et al., 2002; Saha et al., 2002). Further, stimulation of the solitary tract can monosynaptically activate GABAergic NTS cells (Boxwell et al., 2013), suggesting that afferent input can initiate feedforward inhibition. Although input from the gustatory cortex is mainly glutaminergic (Torrealba and Muller, 1996), some cortical input to the NTS makes connections to GABAergic interneurons (Smith and Li, 2000). Temporary pharmacological elimination of GC input to NTS shows similar effects to that reported here: responses to some tastants were attenuated while others were enhanced, sometimes within the same cell (Di Lorenzo and Monroe, 1995). These data suggest that the effects of GABA activation may be at least partially accounted for by mimicking GC input to NTS. Another potential source of GABAergic influence may be the AMG. While anatomical evidence suggests that AMG input to the NTS is inhibitory (Batten et al., 2002; Saha et al., 2002), physiological studies suggest that the effect of stimulation of AMG-NTS input is excitatory (Cho et al., 2003), suggesting the possibility that the AMG generates a disinhibitory effect in the NTS (see Herman et al., 2012). If true, that might contribute to the enhanced responses that became apparent following optical stimulation of GABA release. Since the AMG supplies a rich centrifugal innervation to the rostral NTS (Kang and Lundy, 2009), GABA-induced enhancement of information that discriminates between palatable vs. unpalatable tastants might be mainly due to GABA release from AMG-NTS projections.

### Conclusions

The effects of GABA release during taste stimulation was studied in the NTS of awake, unrestrained rats. GABA changed the taste response profile in about half of the taste responsive cells that were recorded, but the overall interrelationships among the taste-evoked across unit patterns were not altered. Interestingly, GABA activation did not result in more narrow tuning in taste-responsive cells as might have been predicted from studies conducted in anesthetized animals (Smith and Li, 1998). Instead, the population response was essentially remodeled by shifting the identities of the cells conveying specific stimulus-related signals. The coherence of spike activity with the lick pattern was also altered by GABA activation but only in taste-responsive cells. In those cells where GABA activation enhanced taste-related lick coherence information conveyed by temporal coding about taste quality was increased. Most notably, taste-driven GABA activation increased the information conveyed by the temporal characteristics of taste responses about palatability, perhaps in parallel with GABA-induced shifts in lick coherence. In all, this study underscores the fact that GABAergic activation remodel the global population response to taste by both shifts in the responses to taste and the extent to which neural activity reflects licking, in this case. Future experiments should tease apart the effects of the various sources of GABAergic activity to obtain a more precise picture of the role of GABA in the rostral NTS.

## Conflict of Interest

The authors declare no competing financial interests.

## Acknowledgements

Supported by NIDCD Grant RO1-DC006914 to PMD.

